# Neural, physiological and behavioral correlates of empathy for pain in Tourette syndrome

**DOI:** 10.1101/2022.12.19.521056

**Authors:** Ronja Weiblen, Carina Robert, Pauline Petereit, Marcus Heldmann, Thomas F. Münte, Alexander Münchau, Kirsten Müller-Vahl, Ulrike M. Krämer

## Abstract

Persons with Tourette syndrome show altered social behaviors, such as echophenomena and increased personal distress in emotional situations. These symptoms may reflect an overactive mirror neuron system, causing both increased automatic imitation and a stronger tendency to share others’ emotions. To test this, we measured the individual level of echophenomena with a video protocol and experimentally induced empathy for pain in 21 participants with Tourette syndrome and 25 matched controls.

In the empathy for pain paradigm, pictures of hands and feet in painful or neutral situations were presented, while we measured participants’ EEG and skin conductance response. Changes in somatosensory mu suppression during the observation of the pictures and pain ratings were compared between groups and correlations were calculated with the occurrence of echophenomena, self-reported empathy and clinical measures.

Our Tourette syndrome sample showed significantly more echophenomena than controls, but the groups showed no behavioral differences in empathic abilities. However, controls, but not patients with Tourette syndrome, showed the predicted increased mu suppression when watching painful compared to neutral actions. While echophenomena were present in all persons with Tourette syndrome, the hypothesis of an overactive mirror neuron system in Tourette syndrome could not be substantiated. On the contrary, the Tourette syndrome group showed a noticeable lack of mu attenuation in response to pain stimuli. In conclusion, we found a first hint of altered processing of others’ emotional states in a brain region associated with the mirror neuron system.

## Introduction

Gilles de la Tourette Syndrome (TS) is a neuropsychiatric disorder defined by the presence of motor and vocal tics (DSM 5)^1^. Of note however, many TS symptoms are inherently social, socially relevant and/or dependent on the social context. One example are so-called echophenomena, which are defined as the automatic imitation of actions, sounds or language without explicit awareness.^2^ Together with other symptoms such as coprophenomena ^3^ or non-obscene socially inappropriate behaviors (NOSI)^4^, echophenomena indicate that social aspects are central to TS, and that individuals with TS are very sensitive to their social environment ^5^. These observations have instigated research on the specific changes in social cognition in TS, but results so far are inconclusive ^5–11^. For example, individuals with TS reported increased personal distress but decreased perspective taking in the Interpersonal Reactivity Index, a measure of trait empathy ^12^. Based on this and other findings, Eddy ^13^ suggested that TS is characterized by increased automatic sharing of others’ actions and emotions and at the same time reduced mentalizing, possibly reflecting an overactive mirror neuron system (MNS). Similarly, it has been speculated that an overactive MNS might be the cause of echophenomena in TS, as it is critically involved in movement imitation ^2, 14, 15^. The MNS has been implied in many higher order social processes such as action understanding, imitation, perspective taking and empathy^16^, thus providing a framework to explain echophenomena as well as irregularities in social behavior in TS. First neural evidence supports this theory. The temporal parietal junction, which is supposedly part of a top-down control network modulating the MNS^17^, shows irregular activity in TS during social cognition tasks.^10, 18^ Interestingly, TPJ activity covaried with the presence of echophenomena. This supports the idea that control of the MNS might be impaired in TS, leading to its increased activation. Even though the neural basis of changes in social cognition as well as echophenomena might be the same, to our knowledge there are currently no studies investigating social cognition and experimentally measured severity of echophenomena in the same sample. Here, we set out to test the hypothesis that the shared neural basis of those social symptoms of TS is an overactive MNS. We measured behavioral and neural correlates of empathy for pain in TS and related these to the patients’ level of experimentally induced echophenomena.

A well-established neural signature of the human MNS, which is measurable with EEG, is somatosensory mu suppression. Decrease of mu frequency amplitude has been shown in both action observation as well as action execution (for a meta-analysis, see Fox^19^), confirming the mirroring property. Furthermore, increased mu suppression has been linked with empathic abilities in several studies looking at empathy for pain.^20–23^ To test, whether an overactive MNS is the basis of echophenomena as well as altered social behaviors in TS, we thus administered an established empathy for pain paradigm^24^ in a group of TS patients and age-, sex- and education matched controls. Our obtained measures were somatosensory mu suppression as a putative marker for the MNS, the skin conductance response (SCR) as an autonomous response to other’s pain, and behavioral evaluations of the experienced pain as a marker of empathic abilities. The paradigm enables us to measure both empathy, through the manipulation of the observed pain level (painful vs. neutral situation), and perspective taking, by instructing participants to imagine different pain sensitivities of the observed individuals (normal vs. enhanced pain sensitivity). In TS, we expected an increase in behavioral empathy, associated with stronger mu suppression and increased SCR to painful stimuli as neurophysiological markers of increased personal distress. We predicted that deficits in downregulating the automatic sharing of the actorś pain should result in impaired differentiation between onés own and the other’s pain sensitivity in TS. Finally, we investigated the relation between experimentally induced echophenomena and behavioral and neurophysiological measures of empathy. We expected that the hypothesized overactive MNS in individuals with TS would lead to increased mirroring of others’ actions, evident in echophenomena, as well as automatic sharing of emotional states and less distinction between own and other’s perspective, observable as an increase in empathy and decreased perspective taking.

## Materials and methods

### Participants

Data of N = 50 participants were collected from February 2020 to August 2021. This sample size is comparable or larger than similar studies measuring group differences with EEG during behavioral tasks.^25–28^ Of those participants, n = 25 had been diagnosed with TS. They were recruited from the specialized TS outpatient clinic in the Center for Integrative Psychiatry at the University Medical Center Schleswig-Holstein, Campus Lübeck and from the Dept. of Psychiatry, Hannover Medical School, where they received a clinical diagnosis by experienced neurologists (A.M., K. M.-V.). Patients were diagnosed according to DSM-5 criteria.^29^ Based on the collected clinical measures, four TS participants were excluded from all data analyses due to severe depression (n = 2), ongoing substance use disorder (n = 1), and subpar IQ (n = 1). Of the remaining n = 21 participants with TS (17 male, mean age (SD) = 33.0 years (14.0), mean years of education (SD) = 15.7 years (2.7)), four were taking antipsychotics (one each was taking pimozide, olanzapine, aripiprazole and amisulpride) and three were medicated with cannabinoids. Dosages were stable in all patients. Three were left-handed.

Additionally, n = 25 healthy controls (HC) (20 male, mean age (SD) = 33.6 years (13.6), mean years of education (SD) = 15.8 years (2.2)) were individually matched to the TS participants regarding sex, age and years of education. They were recruited by means of poster and online advertisements as well as the university email lists. All controls were free of clinically relevant psychiatric and neurologic symptomatology at the time of testing. One control participant was left-handed. All participants gave their written informed consent to participate in the study in accordance with the Declaration of Helsinki (1964). The local ethics committee gave their approval of the overall project (17-167).

### Clinical, personality and intelligence measures

For matching purposes, we measured general intelligence in all participants with four subtests of the Wechsler Adult Intelligence Scale (WAIS-IV^30^; subtests: similarities, picture completion, arithmetic, coding). We further assessed psychiatric symptomatology with the Mini International Neuropsychiatric Interview (M.I.N.I.). Symptoms of obsessive compulsive disorder (OCD) were scored using the Yale Brown Obsessive Compulsive Scale (YBOCS)^32^ and the Obsessive-Compulsive Inventory (OCI)^33^. Severity of attention deficit hyperactivity disorder (ADHD) symptoms was scored using the ADHD-Index scale of the Conners Adult ADHD Rating Scale (CAARS)^34^. Depressive symptoms were measured with the revised version of the Beck-Depression-Inventory (BDI-II)^35^.

To measure TS symptom severity, several questionnaires were administered on the day of the measurement. Tic severity was assessed using the Yale Global Tic Severity Score (YGTSS)^36^ and the Adult Tic Questionnaire (ATQ)^37^, and premonitory urges using the Premonitory Urge for Tics Scale (PUTS)^38^. Lifetime occurrence of TS symptoms was assessed with the Diagnostic Confidence Index (DCI)^39^. For all participants, standardized video recordings were taken and scored using the Modified Rush Video-Based Tic Rating Scale (MRVS)^40^, which provides a total tic score ranging from 0 to 20.

To assess self-reported empathy in our participants, the Interpersonal Reactivity Index (IRI)^41^ was used. This measure includes affective as well as cognitive aspects of empathy and is divided into four subscales, three measuring aspects of affective (fantasy, empathic concern and personal distress) and one measuring cognitive empathy (perspective taking).

All questionnaires were administered in their German versions. Structured interviews (YGTSS, YBOCS, M.I.N.I, DCI) were done by trained interviewers, the rest of the questionnaires were filled out by the participants either online in the week prior to the measurement or in paper and pencil format on the day of the measurement. TS symptom questionnaires were always administered on the day of the measurement.

### Procedure

Each participant was asked not to take any drugs or alcohol 24 hours prior to the appointment, but exemptions were made for necessary medications. Testing started with the IQ test, followed by two short video recordings for the MRVS (10 min) and the “Echometer” (5 min, explained below). Participants were then prepared for the EEG and SCR measurement and underwent two paradigms, a paradigm investigating provoked speech errors (50 min, unrelated to the present topic of investigation and not included in this paper) and the empathy for pain paradigm (25 min, explained below). The order of the two paradigms was counterbalanced across participants. After the EEG measurement, participants took a break with a duration of their own choosing. Finally, we administered the structured interviews (M.I.N.I, Y-BOCS), all participants filled out the empathy questionnaire (IRI) and TS participants filled out the aforementioned TS symptom questionnaires. Before leaving, participants were debriefed by the experimenters. The whole appointment lasted around five to six hours. Participants were compensated with 15 Euros per hour.

### Echometer

To assess participants’ tendency for echophenomena, we used a similar paradigm as Finis and colleagues.^42^ In the paradigm, the participants observe short video clips while they are simultaneously videotaped, to capture any imitation of movements in the videos. Six different video clips with a duration of three seconds (1 s fixed-image, 1 s tic of a person with TS or spontaneous movement of an actor, 1 s fixed-image) were presented. The video clips showed head and shoulder view segments of one of three individuals with TS (2 male) or one of three actors (2 male). Video clips of the individuals with TS stemmed from previous ten-minute recordings of another study and showed naturally occurring tics. Video clips of the actors were created to match those of the patients and showed spontaneous movements. Each clip included one movement; for spontaneous movements: eyes widening, one eyebrow up and tongue protrusion; for tics: mouth opening, eyebrows up and eyes rolling. Each video was presented five times in a fixed order with a fixation cross (5 s) in between each clip. Thus, each of the 30 trials lasted eight seconds.

Participants were knowingly videotaped using a camera placed on top of the screen. They were instructed to watch the video clips and to fixate on the fixation cross shown in between the video clips. To assure attention, participants were asked to remember which person performed which movement. After the last video clip, the question “Which movement was performed by this person?” was posed, and participants were requested to perform the correct movement. Participants were not instructed to imitate any movements during the task and thus were blinded to the aim of the paradigm.

Videotapes of the participants during the Echometer were rated by two independent raters, who were blinded to the observed videos. We calculated echophenomena frequency for exact repetitions of the observed action (i.e., eye rolling when observing eye rolling) as well as movements in the same body part (i.e., eye movements when observing eye rolling) in the 8 seconds after onset of the video.

### Empathy for pain paradigm

Our empathy for pain paradigm was similar to that of Hoenen and colleagues.^24^ Participants were seated in front of a computer screen and asked to watch 160 pictures. Each picture was presented for 2.5 s followed by a 2.5 s fixation cross (Fig.1). The stimulus set consisted of 80 pictures depicting painful or neutral situations, taken from the Visually induced pain empathy repository (VIPER)^43^. The pictures were presented in four blocks, each consisting of 40 pictures (20/20 painful/neutral). Before each block, a screen with information about the pain sensitivity of the actors in the pictures was provided. In two blocks, participants were told that the actors have a normal pain sensitivity. In the other two blocks, participants were told that the actors have enhanced pain sensitivity, resulting in pain experience even after mild tactile stimulation. The blocks were presented in counterbalanced order and pictures in the blocks were randomized per participant. In 25% of the trials, participants were asked to rate the painfulness of the situation for the actor and for themselves if they were in that situation. Ratings were given on a visual analog scale from 0 = *not at all* to 100 = *very*. All participants rated the same stimuli. Overall, participants watched 80 pictures twice, once in each pain sensitivity condition, resulting in 160 trials. Overall duration of the paradigm was 25 minutes.

**Figure 1.**
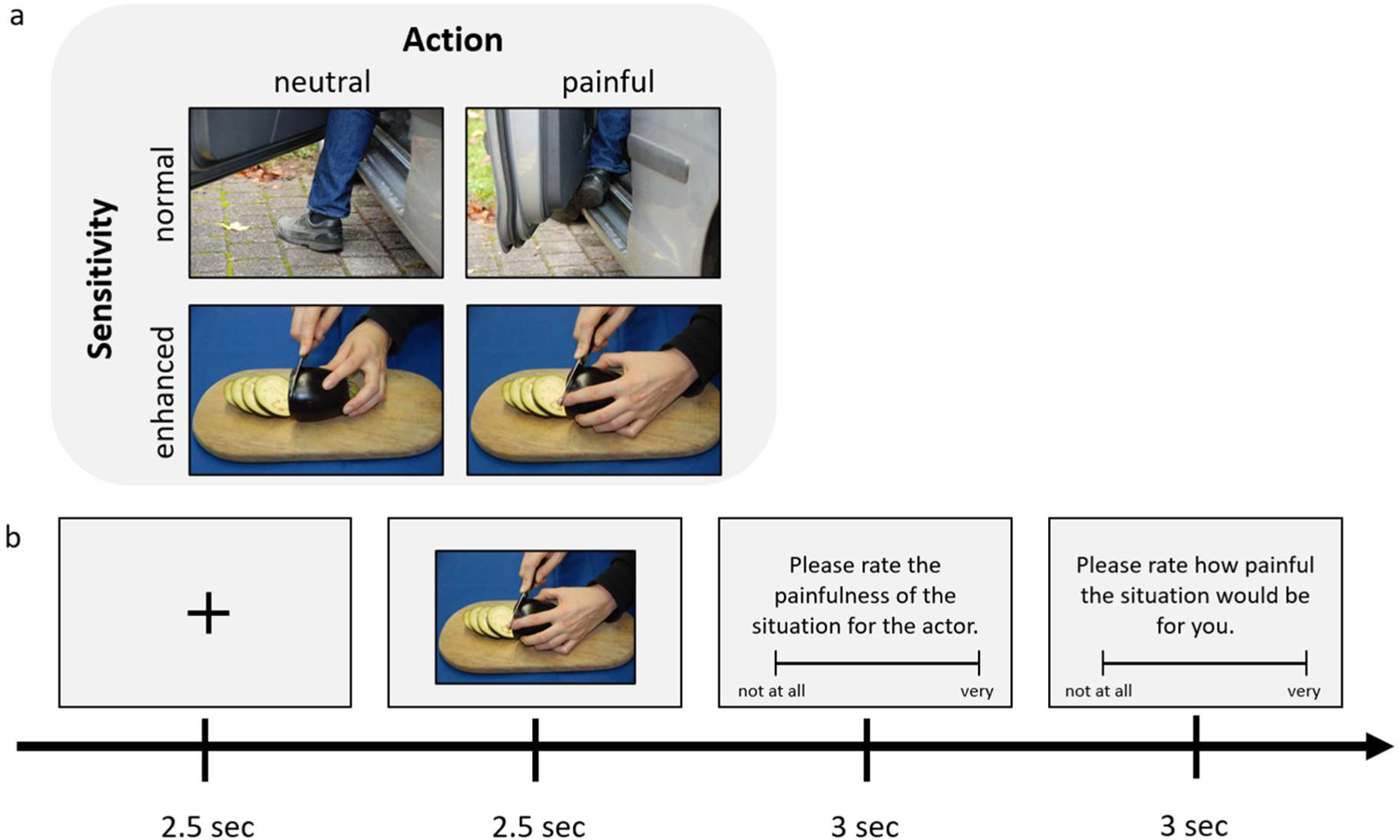
Empathy for pain paradigm. **(a)** Condition matrix with examples of stimuli. *Action* refers to the painfulness of the observed action; *Sensitivity* refers to the imagined pain sensitivity of the observed individual. All stimuli were presented in both sensitivity conditions. **(b)** Trial sequence. Only 25% of the 160 trials were rated.

### Electrophysiological recordings

Electroencephalographic data (EEG) were recorded with 59 Ag/AgCl electrodes placed on an elastic cap according to the international 10–20 system using BrainVision Recorder (BrainProducts GmbH) with the Brainamp MR plus amplifier. An online reference was placed on the left earlobe, and an off-line reference electrode was placed on the right earlobe. Horizontal and vertical EOG was recorded with four electrodes placed next to the outer corners of the eyes and above and below the left eye. Sampling rate was 500 Hz, and data were recorded with an online high-pass filter of 0.016 Hz and a notch filter at 50 Hz. Impedances were kept below 10 kΩ.

Information on skin conductance response (SCR) recording, analysis and results is provided in the supplementary material. It should be noted that the experimental setup, in particular the stimulus timing was not optimized for SCR analyses.

### EEG data analysis

All data preprocessing was done in MATLAB®, using the *eeglab* toolbox^44^. During offline processing, data were down-sampled to 250 Hz, re-referenced to left earlobe and filtered with a high-pass filter at 0.5 Hz and a low-pass filter at 44 Hz. The signal was segmented into epochs from −500 ms to 2500 ms around the beginning of stimulus presentation. Epochs with obvious movement artifacts due to voluntary movements or tics were manually removed as well as channels with very noisy data. Eye movements and further muscle artifacts and channel noise were corrected using independent component analysis (ICA). Visual screening and the automated classifier ICLabel^45^ were used to classify components. On average, 5.9 components were removed per participant. After the ICA, the epoch baseline (−500 0) was removed from each epoch. Then, epochs with remaining artifacts were manually rejected and previously rejected channels were interpolated. On average, 27 epochs were excluded (range: 1 – 86) and 0.7 channels interpolated (range: 0 – 6) per participant. Finally, we calculated the surface Laplacian as suggested by Perrin and colleagues^46^ and implemented in MATLAB® by Cohen^47^.

On average *m* = 28.18 (*sd* = 6.58) trials per combination of conditions remained in the TS group and *m* = 36.99 (*sd* = 2.51) trials in the control group. Participants with more than 50 % of rejected trials or less than 15 trials per combination of conditions were excluded from further analyses. We chose this relatively high threshold, as we had to exclude many trials in the TS group due to movement artifacts because of tics. We had already anticipated this and thus chose a high number of trials to begin with. One participant from the TS group was excluded and none of the healthy controls. Thus, analyses of EEG data are conducted on n = 20 TS patients and n = 25 healthy controls.

On the preprocessed data, we did a time-frequency analysis. We calculated trial-averaged power for each subject, channel and experimental condition for linearly spaced frequencies from 1 Hz to 30 Hz using Morlet wavelet convolution with a variable number of wavelet cycles. Wavelet cycles increased logarithmically from 4 cycles to 15 cycles as a function of frequency. We report decibel (dB) normalized data, using −400 to −100 ms pre stimulus as a baseline.

To check, whether there are group differences in mu power independent of our task manipulations, we looked at absolute and relative mu power in all central clusters, separately for the time of stimulus presentation and the pe-stimulus baseline used previously (−400 to − 100 ms). Absolute power spectral density of the mu band (8-12Hz) was obtained by applying a fast Fourier transform and averaging mu power per cluster and participant. Relative mu power was calculated by dividing mu power spectral density by the absolute power of all frequencies within 1 and 30 Hz.

### Statistical analysis

To compare the two groups regarding demographic measures and clinical symptoms from the questionnaires, we used non-parametric Mann-Whitney-U tests. For group comparisons in the Echometer, we calculated independent t-tests for both measures. For the pain ratings in the empathy for pain paradigm, we calculated a mixed ANOVA. Group was used as a between group factor (TS vs. HC) and action (painful vs. neutral) and sensitivity (normal vs. enhanced) and perspective (otheŕs pain vs. own pain) served as within subject factors. For correlating behavioral results with other measures, we calculated the four difference values for each combination of conditions (e.g., difference of action (painful – neutral) in normal sensitivity condition or difference of sensitivity (enhanced – normal) in painful condition) to get a measure of the magnitude of the action or sensitivity effect in either condition. All described correlations are Pearson’s correlations.

For the statistical analysis of the EEG data during the empathy task, we took the results of the time-frequency analysis and calculated mean power in the mu frequency band (8–12 Hz) for the time window of interest, 500 to 1500ms after stimulus onset. We chose this time window based on the suggestions of Whitmarsh and colleagues^48^, who underlined the importance of not including the time directly after stimulus onset as to not confound the modulation of ongoing alpha activity and stimulus-onset evoked responses. As we were interested in sensorimotor mu suppression, we focused our analysis on a cluster of central electrodes, including C3, C1, CP3, CP1 (left), Cz, CPz (middle) and C2, C4, CP2, CP4 (right). The occipital cluster, including O1, O2 and Oz, served as a control. Power changes were then compared using a mixed ANOVA with group (TS vs. HC) as a between subject factor and action (painful vs. neutral), sensitivity (normal vs. enhanced), region (central vs. occipital) and laterality (right vs. middle vs. left) as within subject factors. Additionally, we calculated Bayesian statistics for the time frequency data, which enabled us to make statements for the null hypothesis. Calculations were done using JASP (Version 0.16.4). We report the results of the Bayesian model averaging (BFincl) implemented as analysis of effects across matched models. We interpret the Bayes factors according to common guidelines, suggesting values > 3 to provide moderate and values > 10 to provide strong support for the alternative hypothesis. Values < 0.33 provide moderate and values < 0.1 strong support for the null hypothesis^49^. For correlating EEG results with other measures, we used the same approach as with the behavioral data and calculated the four difference values for each combination of conditions for mu power in the right central cluster.

Group differences in absolute and relative mu power were tested by means of a mixed ANOVA with group (TS vs. HC) as a between subject factor and region (central right vs. central middle vs. central left) as a within subject factor.

### Data availability

Anonymized data can be shared by request from any qualified investigator. Data will be available for 10 years.

## Results

### Demographics, clinical and questionnaire results

Sex, age, and IQ did not differ between TS patients and healthy controls (Table 1), demonstrating successful matching procedures. TS patients showed significantly more obsessive-compulsive symptoms and symptoms of ADHD. Severity of TS symptoms in our TS group was comparable to similar studies.^5, 15, 50, 51^ Regarding empathic abilities, TS participants reported significantly lower fantasy abilities in the IRI. The two groups did not differ regarding their trait empathic concern, personal distress and perspective taking (Fig. 2a). Additionally, we had hypothesized a correlation between tic severity and trait empathy. Indeed, the YGTSS total tic score correlated with self-reported personal distress in the empathy questionnaire (IRI; *r* = 0.57, *p* =.009), with higher tic severity relating to more reported personal distress (Fig. 2b). Other subscales of the IRI did not significantly correlate with the YGTSS total tic score

**Figure 2.**
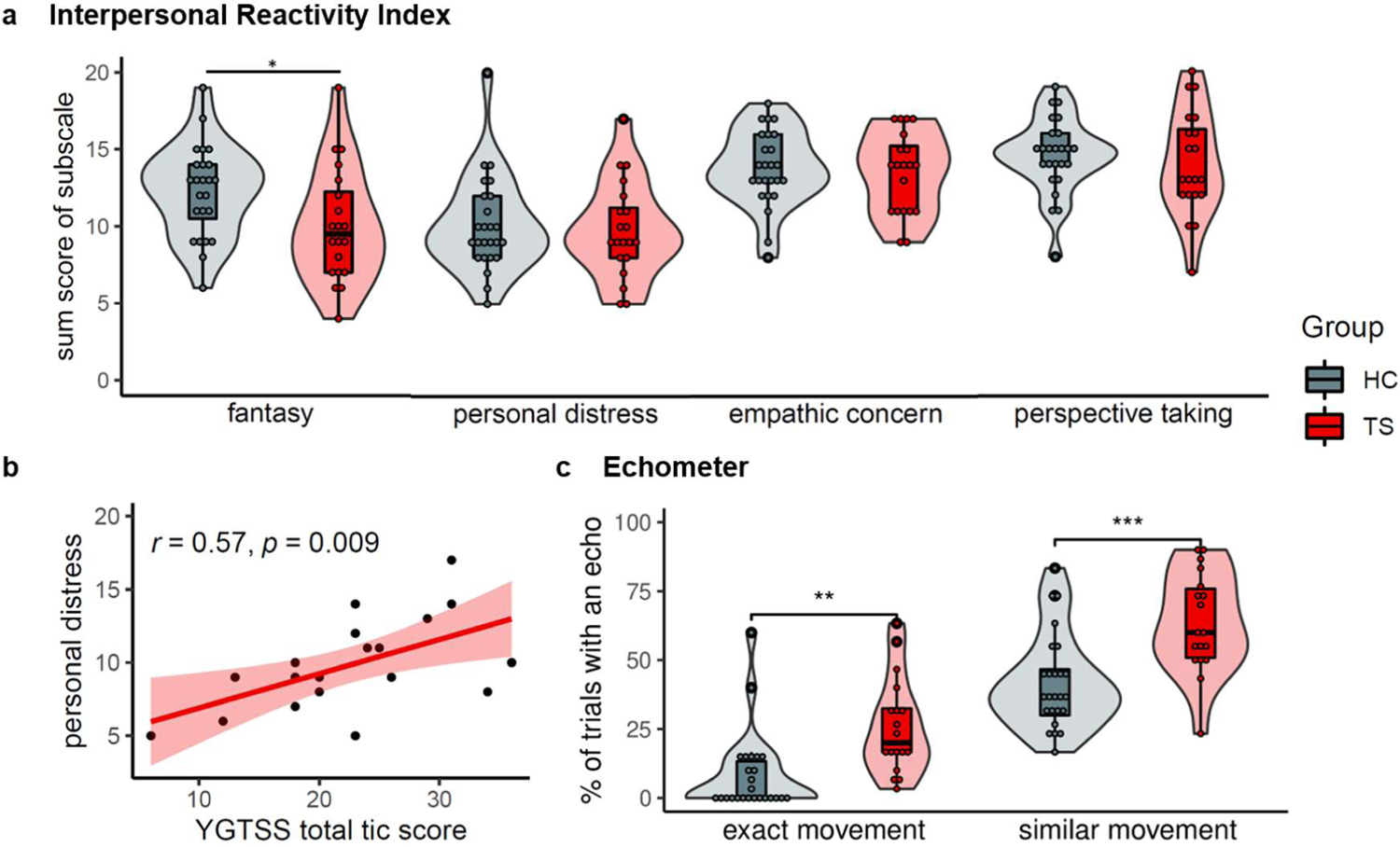
Clinical and behavioral results of n = 25 HC and n = 20 TS. **(a)** Group comparisons with Mann-Whitney U tests of the Interpersonal Reactivity Index (IRI). Significant group difference for subscale fantasy: *U* = 331.5, *p* = .031; but not personal distress: *U* = 260, *p* = .826; empathic concern: *U* = 262, *p* = .791; perspective taking: *U* = 281, *p* = .483. **(b)** Correlation in TS group of the subscale personal distress in the IRI and total tic score as measured by the Yale Global Tic Severity Scale (YGTSS). **(c)** Number of echoes in Echometer. Group comparisons with t-tests revealed significant differences for both exact replications: *t*(1,32.58) = −3.49, *p* = .001; as well as movements in the same body parts: *t*(1,35.58) = −3.98, *p* < .001.

**Table 1.** Overview of personality and clinical measures.

| Questionnaire | TS | HC | N | U | p | r |
| --- | --- | --- | --- | --- | --- | --- |
| **Sex** | 17 m, 4 f | 20 m, 5 f |  |  |  |  |
| **Age** | 27 (23) | 30 (27) | 21/25 | 272.5 | .834 | .03 |
| **IQ** | 104 (13) | 102 (14) | 21/25 | 279.5 | .715 | .06 |
| **Obsessive-compulsive behavior** |  |  |  |  |  |  |
| YBOCS thoughts | 1 (6) | 0 (0) | 21/25 | 133 | < .001 | .55 |
| YBOCS actions | 5 (7) | 0 (0) | 21/25 | 107.5 | < .001 | .62 |
| YBOCS total | 8 (12) | 0 (0) | 21/25 | 96.5 | < .001 | .65 |
| OCI | 16 (25) | 7 (9) | 21/25 | 148 | .012 | .37 |
| **Depression** | 8 (15) | 3 (7) | 21/25 | 177 | .060 | .28 |
| **ADHD** | 11 (11) | 6 (7) | 21/25 | 154 | .017 | .35 |
| **Empathy (IRI)** |  |  |  |  |  |  |
| Fantasy | 9.5 (5.25) | 13 (3.5) | 20/25 | 331.5 | .031 | .33 |
| Empathic Concern | 14 (4.25) | 14 (3) | 20/25 | 262 | .791 | .04 |
| Personal Distress | 9 (3.25) | 9 (4) | 20/25 | 260 | .826 | .03 |
| Perspective Taking | 13 (4.25) | 15 (2) | 20/25 | 281 | .483 | .11 |
| **TS Symptomatology** |  |  |  |  |  |  |
| MRVS total score | 12 (8) | 3 (4) | 21/25 | 10 | < .001 | .83 |
| YGTSS Total motor tic score | 14 (2) |  |  |  |  |  |
| YGTSS Total vocal tic score | 8 (7) |  |  |  |  |  |
| YGTSS Total tic score | 23 (8) |  |  |  |  |  |
| YGTSS Global severity score | 42 (15.25) |  |  |  |  |  |
| ATQ | 31 (19) |  |  |  |  |  |
| PUTS | 21 (4) |  |  |  |  |  |
| DCI | 58 (18) |  |  |  |  |  |

Note. Questionnaire data separated by group. Median scores plus interquartile range in brackets. We report results of a Mann Whitney U test to compare the groups. m = male, f = female. IQ, Intelligence Quotient as measured by subtests of the Wechsler Adult Intelligence Scale (WAIS-IV). YBOCS, Yale Obsessive Compulsive Inventory. OCI, Obsessive Compulsive Inventory. ADHD, Attention Deficit Hyperactivity Disorder as measured by the Conners Adult ADHD Rating Scale. IRI, Interpersonal Reactivity Index. MRVS, Modified Rush Video-Based Tic Rating Scale. YGTSS, Yale Global Tic Severity Scale. ATQ, Adult Tic Questionnaire. PUTS, Premonitory Urges for Tics Scale. DCI, Diagnostic Confidence Index. TS, Tourette Syndrome. HC, Healthy Controls.

### Behavioral task results

The analysis of echophenomena yielded clear group differences for both exact imitations (*t*(1,32.58) = −3.49, *p* = .001, *d* = −1.11) as well as movements in the same body part (*t*(1,35.58) = −3.98, *p* < .001, *d* = −1.24). As expected, participants of the TS group showed a higher tendency to echo what they observe than the healthy controls (Fig. 2c).

In the empathy for pain paradigm, we found an interaction effect of action, sensitivity and perspective (*F*(1,44) = 11.25, *p* = .002, *η*^2^ < .01). We followed this up with separate ANOVAS for each perspective and found an interaction of action and sensitivity on ratings of others’ pain (*F*(1,44) = 21.69, *p* < .001, *η*^2^ = .02), with all four Bonferroni corrected pairwise comparisons being significant. When looking at the ratings of the pain that participants themselves would feel in the observed situation, we again found an interaction of action and sensitivity (*F*(1,44) = 4.16, *p* = .047, *η*^2^ < .01). However, Bonferroni corrected pairwise comparisons revealed no differences between sensitivity conditions in both painful and neutral stimuli. This shows that both HC and TS performed as expected: They rated the other’s pain higher in the painful condition compared to the neutral condition; and they rated the other’s pain higher in the enhanced pain sensitivity condition compared to the normal pain sensitivity condition. When rating their own imagined pain, participants made no difference between sensitivity conditions but rated painful stimuli as more painful than neutral ones. However, we did not find any group effects in the ratings. The three-way interaction of group, action and sensitivity on the rating of others’ pain was not significant (*F*(1,44) = 0.13, *p* = .72, *η*^2^ < .01, Fig. 3), although the differences between sensitivity conditions in the TS group were nominally smaller. On the behavioral level, the experimental manipulations were thus successful, but none of the hypothesized group differences were confirmed. Detailed results of the ANOVA and post hoc tests can be found in the supplementary materials.

**Figure 3.**
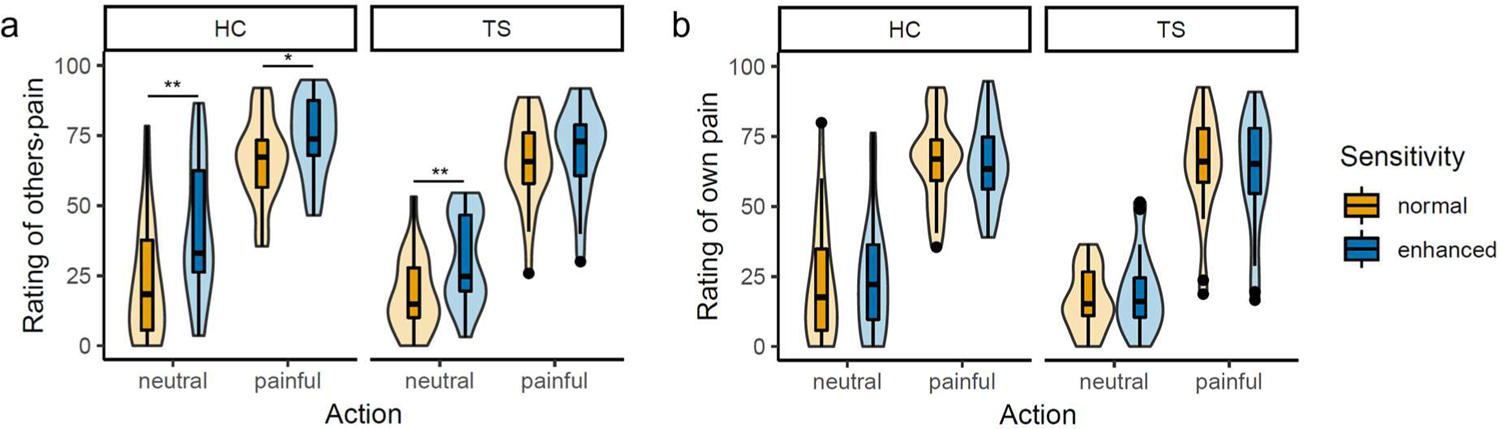
Pain ratings in empathy for pain paradigm. A mixed Anova revealed a significant interaction of action, sensitivity and perspective (*F*(1,44) = 11.25, *p* = .002) for n = 20 TS and n = 25 HC. **(a)** Rating of painfulness for the actor in the picture from „not at all” to „very” under the normal (yellow) and enhanced (blue) sensitivity condition for both healthy controls (HC, left side) and Tourette patients (TS, right side). **(b)** Rating of own imagined pain in the depicted situation on same scale.

### Electrophysiological results

The mu suppression in the empathy for pain paradigm showed an interaction of all factors (group, action, sensitivity, region & laterality): *F*(2,82) = 3.37, *p* = .039, *η*^2^ < .01 (Fig. 4). To understand this interaction, we calculated ANOVAs separately for central and occipital sites. At occipital electrodes, we found a main effect of action (*F*(1,43) = 13.80, *p* = .001, *η*^2^ = .01), indicating more pronounced alpha suppression after observing painful compared to neutral actions. At the central electrodes, we found a significant interaction of group, action and sensitivity (*F*(1,41) = 6.52, *p* = .014, *η*^2^ < .01, BF = 4.58). Following ANOVAs for each of the groups revealed no significant interaction of action and sensitivity in the TS group (*F*(1,17) = 2.24, *p* = .153, *η*^2^ < .01, BF = 2.62), nor any other effects. In HC we found an interaction of action and sensitivity (*F*(1,24) = 5.11, *p* = .033, *η*^2^ < .01, BF = 2.26). HC showed a simple main effect of action in the normal sensitivity condition (*F*(1) = 5.34, *p* = .03) but not in the enhanced sensitivity condition (*F*(1) = 0.03, *p* = .86). Finally, there were no group differences in absolute or relative mu power, neither during the pre-stimulus baseline period (absolute: *F*(1,43) = 0.07, *p* = .790, *η*^2^ < .01; relative: *F*(1,43) = 0.14, *p* = .707, *η*^2^ < .01) nor during the stimulus presentation (absolute: *F*(1,43) = 1.98, *p* = .166, *η*^2^ = .03; relative: *F*(1,43) = 0.76, *p* = .387, *η*^2^ = .02).

**Figure 4.**
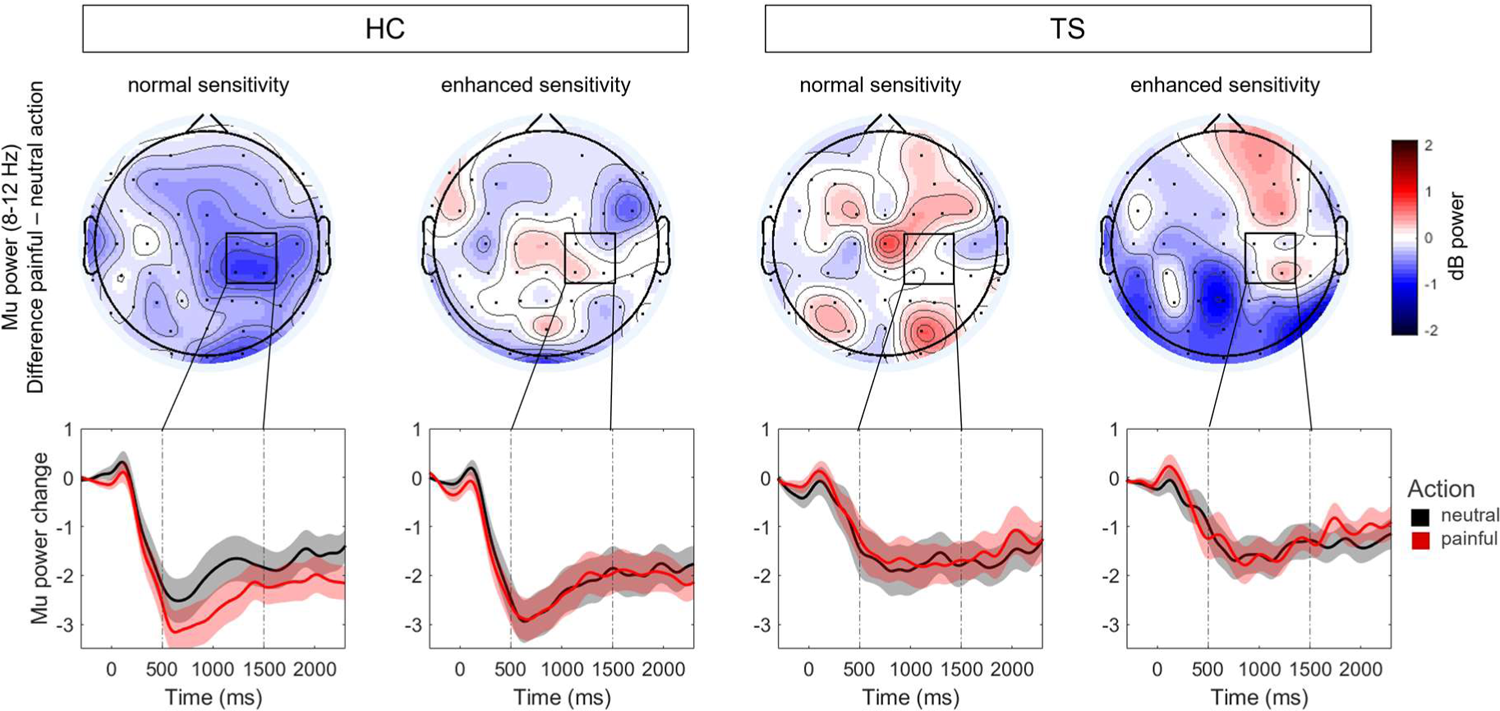
Baseline corrected mu power (8 – 12 Hz) in empathy for pain paradigm. **(Top row)** Spatial distribution of mu power differences (painful-neutral) in time-window of interest (500 – 1500 ms). **(Bottom row)** Time course of mu power in right central cluster for the painful (red) and neutral (black) condition, separately for n = 25 healthy controls (HC, left columns) and n = 20 Tourette patients (TS, right columns) and for normal and enhanced sensitivity. Time-point 0 refers to the picture onset.

### Correlations of echophenomena with empathy measures

Based on our hypothesis on a shared mechanism behind echophenomena and empathy, we expected the number of echophenomena to correlate with our behavioral, physiological and neural indices of empathic abilities in the whole sample. To test this, we focused on the difference of action (painful – neutral) in the normal sensitivity condition. However, echophenomena did not correlate with trait personal distress (*r* = 0.22, *p* = .15), behavioral (*r* = 0.27, *p* = .08), physiological (SCR: *r* = 0.11, *p* = .50) or neural empathy measures (mu: *r* = 0.15, *p* = .35). Additionally, we hypothesized a correlation between tic severity and empathy in the TS group. However, no correlations could be found between tic severity measures of the YGTSS or the MRVS video protocol with behavioral, neural or physiological empathy measures.

## Discussion

In this study, we tested the hypothesis that echophenomena in TS are related to increased mirror neuron system activity and increased automatic sharing of others’ emotional states. As expected, we found that participants with TS showed more echophenomena than controls. However, we did not find any empathy related behavioral differences between TS and HC and no differences in trait empathy or skin conductance between our groups. Interestingly, contrary to our hypothesis, the TS group showed reduced pain-related mu suppression, indicating altered processing of others’ emotional states in a brain region associated with the MNS.

### Echophenomena and trait empathy in TS

In line with a previous study^42^, we found a strong difference between TS and controls in the frequency of echophenomena induced by our Echometer paradigm. This was true for both exact replications of the observed movement as well as movements with the same body part. Furthermore, all participants with TS showed echoes, which stresses the high prevalence of echophenomena in this patient group. This is in contrast to previous, lower estimates of its prevalence, usually ranging below fifty percent.^52, 53^ The difference is most likely due to the mode of measurement, as prevalence estimates in these studies were based on self-report or clinicians’ observations. It is thus important for future explorations of the neural basis of echophenomena to experimentally measure their occurrence. Additionally, echoes were not only seen in the TS patient group but also, with much lower frequency, in the healthy controls, underlining that imitating others is a normal process and not per se pathological.^2^ In conclusion, our TS sample exhibited the expected increased tendency for echophenomena compared to controls and should thus, according to our hypothesis, show further symptoms of altered social behaviors and an overactive MNS.

One of the previous findings which had prompted our hypothesis was the report of increased self-reported personal distress, combined with reduced perspective-taking in the Interpersonal Reactivity Index (IRI) in a relatively large TS sample.^12^ The IRI personal distress subscale measures unease and anxiety in negative emotional settings and is related to poor social functioning, whereas the perspective taking subscale measures the tendency to think about the otheŕs point of view and is correlated with higher social functioning.^54^ We utilized the same questionnaire but could not replicate the results in our sample. TS participants in our study did not report increased personal distress or reduced perspective taking compared to the controls. We did find an increase of reported personal distress with higher tic severity, but this was also contrary to what Eddy and colleagues reported^12^, as they found a *negative* correlation between tics and personal distress. Although our sample was smaller compared to theirs, our effect sizes were also considerably smaller, which makes it unlikely that we missed the effect only because of a smaller sample. Furthermore, our sample included far less medicated participants, but comorbidities were comparable across both samples. TS participants in our study did report lower fantasy scores compared to controls, indicating a decreased tendency to imaginatively transpose themselves into fictional situations. Overall, we found group differences in self-reported empathy but those were not consistent with the previous report of Eddy and colleagues.^12^ Additionally, the results of the empathy self-report did not correlate with the tendency for echophenomena.

### Empathy for pain in TS

In the empathy for pain paradigm, both groups performed similarly and in line with the task manipulation, showing that task instructions were understood by both groups. When rating their own supposed pain, both groups performed similarly, suggesting no difference in their perception of the painfulness of the stimuli. We had hypothesized increased painfulness ratings after painful stimuli for our TS group, which we could not confirm. However, one could argue that the pain ratings in our paradigm did not reflect only automatic sharing of the experience of the actor but also cognitive understanding of the situation. Furthermore, the pain stimuli were not overly graphic, did not show obvious signs of injuries, painful facial expressions or body language and the instruction referred to actors, underlining that the pictures were not real. The stimuli might thus not elicit strong discomfort in the viewer. Overall, our paradigm might not have been complex enough to detect subtle behavioral differences between the groups. Previous studies reporting group differences in social cognition used different stimuli, such as emotional facial expressions, faux pas stories or humorous cartoons, which might be more cognitively demanding and thus measuring different aspects of empathy and perspective taking.^5–11^ Moreover, TS might still show altered neural processing even in the absence of obvious behavioral differences.

In terms of the neural data, several studies have probed mu suppression under different conditions or in different groups with an almost identical setup as ours: participants observing pictures of body parts in painful or neutral situations.^24, 55–57^ Those studies consistently find increased mu suppression during the observation of painful compared to neutral pictures. The magnitude of mu suppression is interpreted as a measure of empathy and is modulated by several factors, with less mu suppression being observed in men than women^57^, during negative mood compared to positive or neutral mood^56^ and after reported parental emotional invalidation.^55^ In line with those findings, our healthy controls showed the expected pain-related mu suppression over the right sensorimotor cortex. This was modulated by our additional manipulation of the actors’ pain sensitivity. Suppression only differed between painful versus neutral stimuli when the pain sensitivity of the actor was normal. This effect vanished under the enhanced pain sensitivity condition, indicating that controls processed both kinds of stimuli as painful in this condition. Our results thus mirror those of Hoenen and colleagues^24^, who originally developed this paradigm and found increased mu suppression in trials where participants were informed that the actors have an increased pain sensitivity. In a similar setup, participants observed hands either being pricked by a needle or being touched by a Q-tip.^23^ Participants were asked to assume either normal pain reactions or switched pain reactions, where needle pricks were not painful, but Q-tip touches were. When participants assumed normal pain reactions, the classic increase of mu suppression in painful over neutral situations was observed. However, when participants assumed switched pain reactions, mu suppression was equally large in painful and neutral situations. Those results combined with what we observed in the healthy controls indicate that mu suppression is not only elicited by automatic empathic reactions to the observed stimuli but can be modulated by top-down processes, making it possible to feel empathy for pain even in situations that would not be painful for the participants themselves. Interestingly, TS participants did not show any significant pain-related attenuation of mu power, although Bayesian analyses indicated equivocal evidence for and against this effect. However, this group difference in pain-related mu suppression speaks against our hypothesis of increased automatic sharing in TS, but rather points in the opposite direction. One might argue that a generally overactive MNS in TS could lead to less pronounced mu attenuation during our paradigm. However, the TS group showed similar absolute and relative mu power during the baseline as well as averaged across all stimulus presentations. This questions that the group differences can be attributed to baseline differences and a generally overactive MNS.

### Is an overactive MNS in TS causing echophenomena and changes in empathy?

What are the implications for our hypothesis that an overactive MNS is the basis of both altered social behaviors and echophenomena? First of all, we did not find any alterations in empathic behavior in the TS group and can thus not draw any conclusion on possible causes of altered social behavior in TS. Second, we did not find evidence that increased somatosensory mu suppression is at the core of the clearly increased tendency for echophenomena in our TS sample. On the contrary, we found less pain-related mu attenuation in the TS group during the empathy for pain task.

Our results might thus support an alternative hypothesis about the basis of echophenomena. Rather than echophenomena being indicative of a generally increased tendency to imitate, they might reflect the failure to inhibit imitation in contexts where it is not useful.^2^ Thus, in situations where imitation has a social function and can be found in healthy adults, we find no increase of imitation in TS compared to controls. It is in situations where imitation serves no purpose, such as our Echometer paradigm, that we find increased imitation in TS. This also fits with previous results in an automatic imitation paradigm.^15^ Both TS and HC showed a comparable modulation of reaction times by incongruent movements compared to congruent movements, indicating similar automatic imitation tendencies. In conclusion, in social contexts where it is beneficial, such as an empathy paradigm, imitation or automatic sharing of the otheŕs state does not seem to be increased in TS compared to controls. This interpretation is further supported by the fact that we found no relationship between the extent of echophenomena or tic severity and the behavioral and neural results in the empathy for pain paradigm.

However, not only did we not find an increase in automatic sharing in our empathy paradigm, our results even pointed in the opposite direction, with TS participants showing less pain-related mu suppression than controls. This fits to an additional hypothesis brought forward by Ganos and colleagues.^2^ They propose that people with TS compensate the deficient inhibition in motor networks with increased activity in inhibitory circuits outside the motor network, presumably prefrontal areas. They thus assume that compensatory activity in the TS brain leads to less responsiveness to experimental manipulations. It is feasible that due to the failure to inhibit tics or echophenomena in inappropriate contexts, the TS brain overcompensates by generally decreasing the responsiveness to stimuli. This fits to recent Go/No-Go studies in TS, which found reduced task-related activations compared to healthy controls.^58, 59^ In one study, task-related BOLD signal change was decreased in cortical motor areas but not in other areas in people with TS. This was accompanied by poorer task performance compared to healthy controls.^59^ In another Go/No-Go task, participants with TS showed less beta suppression in the sensorimotor area but enhanced beta power in parieto-occipital brain regions compared to healthy controls, but task performance was the same in both groups.^58^ Similarly, in the aforementioned automatic imitation task^15^, where TS and HC showed no differences in the interference effect, people with TS exhibited less movement facilitation by observing congruent biological movements than healthy controls. However, they showed more interference than healthy controls when observing incongruent biological movements compared to the observation of non-biological movement, hinting to an overactive imitation inhibition system that is active during the task. To conclude, our findings of decreased mu modulation in TS compared to the controls might be explained by increased inhibition. However, it is important to note, that none of these studies or our study provided direct neural evidence for increased inhibitory activity.

### Limitations

Our groups were of moderate size, which might have led us to miss small group differences. The results of Bayes statistics underlined this problem, as we found moderate evidence for group differences in pain-related mu suppression in the interaction, but overall weak effects of the pain modulation in both groups. The small magnitude of pain-related mu suppression might be partly due to our sample being predominantly male, as male participants were found to exhibit less mu suppression than females in previous research.^57^ Also the number of trials in each experimental condition was kept relatively low to reduce fatigue during the task, but might have been too low to detect rather subtle condition or group differences. Furthermore, trials with tics had to be removed due to movement artifacts which might have led us to miss effects during tics. Another caveat regarding our results is the heterogeneity of our TS sample. TS is often accompanied by several comorbidities, and this was the same in our group. Compared to the controls, the TS group showed more symptoms of depression, anxiety, ADHD and OCD, all of which could have influenced our results. However, as most individuals with TS suffer from symptoms of one or more of these comorbidities, we considered it best to include them to create a representative sample for this clinical group. Furthermore, while some previous studies looking at social cognition differences in TS only included uncomplicated TS^7, 10^, most included comorbid ADHD and/or OCD^5, 9, 12^. Furthermore, our empathy paradigm was relatively easy and might thus not be suitable to measure subtle group differences in behavioral empathy. Finally, previous studies, which reported differential activity in brain regions associated with the MNS focused on the temporoparietal junction ^10,18^. As we focused on sensorimotor activity, we cannot rule out altered activity in other regions of the MNS in individuals with TS based on our data.

### Conclusions

Together, our results could not support the theory of an overactive MNS based on examination of the sensorimotor cortex, and we found no evidence of increased levels of empathy in TS. In contrast, our findings indicate reduced pain-related neural processing in TS, In contrast, our findings indicate reduced pain-related neural processing in TS, although this had no behavioral impact in the current task. As our TS sample evinced high levels of echophenomena, the underlying neurocognitive processes of this symptom remain to be established. Overall, as our study, as well as previous studies^10, 18, 60^ imply altered processing of stimuli in social contexts, future research should continue to explore the neural basis of social symptoms and echophenomena in TS.

## Supplementary material

Supplementary material is available online.

## Abbreviations

HC: healthy control
MNS: mirror neuron system
SCR: skin conductance response
TS: Tourette syndrome

