## Supplementary Materials for "Neural, physiological and behavioral correlates of empathy for pain in Tourette syndrome"

#### Abbreviations & Factors Overview

ges = generalized eta squared

TS = Tourette Syndrome

HC = Healthy Controls

Explanation of factors:

- Group: TS & HC
- Action: painful & neutral (pictures shown during paradigm)
- Sensitivity: normal & enhanced (instructed pain sensitivity of actor in pictures)
- Perspective: own pain & other pain (whose pain is rated after the picture?)
- Region: central & occipital (electrode location for EEG)
- Laterality: middle & left & right (electrode location for EEG)

#### Behavioral Results Empathy for Pain Task

**Table S1**: Summary statistics of pain ratings per combination of conditions:

| Group | Action | Sensitivity | Perspective | n | mean | sd |
|---|---|---|---|---|---|---|
| HC | neutral | normal | Own pain | 25 | 22.51 | 21.09 |
| HC | neutral | enhanced | Own pain | 25 | 23.88 | 20.18 |
| HC | painful | normal | Own pain | 25 | 66.52 | 15.14 |
| HC | painful | enhanced | Own pain | 25 | 65.50 | 14.69 |
| TS | neutral | normal | Own pain | 21 | 16.39 | 10.87 |
| TS | neutral | enhanced | Own pain | 21 | 19.09 | 14.07 |
| TS | painful | normal | Own pain | 21 | 64.43 | 18.92 |
| TS | painful | enhanced | Own pain | 21 | 61.57 | 20.88 |
| HC | neutral | normal | Other pain | 25 | 22.74 | 21.14 |
| HC | neutral | enhanced | Other pain | 25 | 41.66 | 23.95 |
| HC | painful | normal | Other pain | 25 | 66.39 | 14.92 |
| HC | painful | enhanced | Other pain | 25 | 74.63 | 13.92 |
| TS | neutral | normal | Other pain | 21 | 18.30 | 13.60 |
| TS | neutral | enhanced | Other pain | 21 | 31.35 | 16.03 |
| TS | painful | normal | Other pain | 21 | 65.71 | 15.66 |
| TS | painful | enhanced | Other pain | 21 | 69.62 | 16.17 |

**Table S2**: Anova with factors Group, Action, Sensitivity and Perspective:

| Effect | DFn | DFd | F | p | p<.05 | ges |
|---|---|---|---|---|---|---|
| Group | 1 | 44 | 1.696 | 0.200 | | 0.02 |
| Action | 1 | 44 | 203.106 | 0.000 | * | 0.61 |
| Sensitivity | 1 | 44 | 18.799 | 0.000 | * | 0.03 |
| Perspective | 1 | 44 | 29.732 | 0.000 | * | 0.03 |
| Group:Action | 1 | 44 | 0.345 | 0.560 | | 0.00 |
| Group:Sensitivity | 1 | 44 | 1.098 | 0.300 | | 0.00 |
| Group:Perspective | 1 | 44 | 0.143 | 0.707 | | 0.00 |
| Action:Sensitivity | 1 | 44 | 14.255 | 0.000 | * | 0.01 |
| Action:Perspective | 1 | 44 | 11.69 | 0.001 | * | 0.00 |
| Sensitivity:Perspective | 1 | 44 | 47.013 | 0.000 | * | 0.03 |
| Group:Action:Sensitivity | 1 | 44 | 0.049 | 0.826 | | 0.00 |
| Group:Action:Perspective | 1 | 44 | 1.057 | 0.310 | | 0.00 |
| Group:Sensitivity:Perspective | 1 | 44 | 2.292 | 0.137 | | 0.00 |
| Action:Sensitivity:Perspective | 1 | 44 | 11.25 | 0.002 | * | 0.00 |
| Group:Action:Sensitivity:Perspective | 1 | 44 | 1.764 | 0.191 | | 0.00 |

**Table S3:** Anova for ratings of other's pain:

| Effect | DFn | DFd | F | p | p<.05 | ges |
|---|---|---|---|---|---|---|
| Group | 1 | 44 | 1.95 | 0.170 | | 0.02 |
| Action | 1 | 44 | 187.46 | 0.000 | * | 0.58 |
| Sensitivity | 1 | 44 | 36.83 | 0.000 | * | 0.09 |
| Group:Action | 1 | 44 | 0.58 | 0.449 | | 0.00 |
| Group:Sensitivity | 1 | 44 | 1.97 | 0.167 | | 0.01 |
| Action:Sensitivity | 1 | 44 | 21.69 | 0.000 | * | 0.02 |
| Group:Action:Sensitivity | 1 | 44 | 0.13 | 0.720 | | 0.00 |

**Table S4**: Pairwise Comparisons *Group* per Action and Sensitivity:

| Action | Sensitivity | variable | group1 | group2 | n1 | n2 | p |
|---|---|---|---|---|---|---|---|
| neutral | normal | Other pain | HC | TS | 25 | 21 | 0.412 |
| neutral | enhanced | Other pain | HC | TS | 25 | 21 | 0.100 |
| painful | normal | Other pain | HC | TS | 25 | 21 | 0.882 |
| painful | enhanced | Other pain | HC | TS | 25 | 21 | 0.265 |

**Table S5**: Pairwise Comparisons *Action* per Sensitivity:

| Sensitivity | variable | group1 | group2 | n1 | n2 | p |
|---|---|---|---|---|---|---|
| normal | Other pain | neutral | painful | 46 | 46 | 0.000 |
| enhanced | Other pain | neutral | painful | 46 | 46 | 0.000 |

**Table S6:** Pairwise Comparisons *Sensitivity* per Action:

| Action | variable | group1 | group2 | n1 | n2 | p |
|---|---|---|---|---|---|---|
| neutral | Other pain | normal | enhanced | 46 | 46 | 0.000 |
| painful | Other pain | normal | enhanced | 46 | 46 | 0.049 |

**Table S7**: Anova for ratings of own supposed pain:

| Effect | DFn | DFd | F | p | p<.05 | ges |
|---|---|---|---|---|---|---|
| Group | 1 | 44 | 1.19 | 0.281 | | 0.02 |
| Action | 1 | 44 | 207.30 | 0.000 | * | 0.62 |
| Sensitivity | 1 | 44 | 0.00 | 0.965 | | 0.00 |
| Group:Action | 1 | 44 | 0.16 | 0.691 | | 0.00 |
| Group:Sensitivity | 1 | 44 | 0.01 | 0.910 | | 0.00 |
| Action:Sensitivity | 1 | 44 | 4.16 | 0.047 | * | 0.00 |
| Group:Action:Sensitivity | 1 | 44 | 0.66 | 0.422 | | 0.00 |

**Table S8**: Pairwise Comparisons *Sensitivity* per Group and Action:

| Group | Action | variable | group1 | group2 | n1 | n2 | p |
|---|---|---|---|---|---|---|---|
| HC | neutral | Own pain | normal | enhanced | 25 | 25 | 0.815 |
| HC | painful | Own pain | normal | enhanced | 25 | 25 | 0.810 |
| TS | neutral | Own pain | normal | enhanced | 21 | 21 | 0.490 |
| TS | painful | Own pain | normal | enhanced | 21 | 21 | 0.645 |

### EEG Results in Empathy for Pain Task

**Table S9:** Summary statistics for mu/alpha power:

| Group | Sensitivity | Action | Region | Laterality | n | mean | sd |
|---|---|---|---|---|---|---|---|
| HC | normal | neutral | central | left | 25 | -2.22 | 1.74 |
| HC | normal | neutral | central | middle | 25 | -1.99 | 1.74 |
| HC | normal | neutral | central | right | 25 | -2.03 | 2.17 |
| HC | enhanced | neutral | central | left | 25 | -2.44 | 2.22 |
| HC | enhanced | neutral | central | middle | 25 | -2.43 | 2.01 |
| HC | enhanced | neutral | central | right | 25 | -2.43 | 2.21 |
| HC | normal | painful | central | left | 25 | -2.40 | 2.14 |
| HC | normal | painful | central | middle | 25 | -2.50 | 2.13 |
| HC | normal | painful | central | right | 25 | -2.71 | 2.03 |
| HC | enhanced | painful | central | left | 25 | -2.52 | 1.96 |
| HC | enhanced | painful | central | middle | 25 | -2.26 | 1.85 |
| HC | enhanced | painful | central | right | 25 | -2.42 | 2.12 |
| TS | normal | neutral | central | left | 19 | -1.64 | 1.66 |
| TS | normal | neutral | central | middle | 19 | -1.57 | 1.50 |
| TS | normal | neutral | central | right | 20 | -1.74 | 2.09 |
| TS | enhanced | neutral | central | left | 20 | -1.63 | 1.90 |
| TS | enhanced | neutral | central | middle | 20 | -1.49 | 1.95 |
| TS | enhanced | neutral | central | right | 20 | -1.49 | 1.59 |
| TS | normal | painful | central | left | 20 | -2.08 | 2.00 |
| TS | normal | painful | central | middle | 19 | -1.55 | 1.62 |
| TS | normal | painful | central | right | 20 | -1.69 | 1.79 |
| TS | enhanced | painful | central | left | 20 | -1.99 | 1.83 |
| TS | enhanced | painful | central | middle | 20 | -1.96 | 2.05 |
| TS | enhanced | painful | central | right | 20 | -1.49 | 1.41 |
| HC | normal | neutral | occipital | left | 25 | -2.85 | 2.07 |
| HC | normal | neutral | occipital | middle | 25 | -2.02 | 2.39 |
| HC | normal | neutral | occipital | right | 25 | -2.76 | 2.58 |
| HC | enhanced | neutral | occipital | left | 25 | -2.85 | 2.36 |
| HC | enhanced | neutral | occipital | middle | 25 | -2.19 | 2.82 |
| HC | enhanced | neutral | occipital | right | 25 | -3.07 | 3.16 |
| HC | normal | painful | occipital | left | 25 | -3.03 | 2.38 |
| HC | normal | painful | occipital | middle | 25 | -2.64 | 2.61 |
| HC | normal | painful | occipital | right | 25 | -3.51 | 2.67 |
| HC | enhanced | painful | occipital | left | 25 | -3.26 | 2.48 |
| HC | enhanced | painful | occipital | middle | 25 | -2.87 | 2.46 |
| HC | enhanced | painful | occipital | right | 25 | -3.44 | 2.88 |
| TS | normal | neutral | occipital | left | 20 | -3.10 | 2.66 |
| TS | normal | neutral | occipital | middle | 20 | -2.30 | 2.47 |
| TS | normal | neutral | occipital | right | 20 | -3.73 | 2.84 |
| TS | enhanced | neutral | occipital | left | 20 | -2.86 | 2.25 |
| TS | enhanced | neutral | occipital | middle | 20 | -1.79 | 2.21 |
| TS | enhanced | neutral | occipital | right | 20 | -2.76 | 2.59 |
| TS | normal | painful | occipital | left | 20 | -3.27 | 2.60 |

| | | | | | | | |
|---|---|---|---|---|---|---|---|
| TS | normal | painful | occipital | middle | 20 | -2.49 | 2.04 |
| TS | normal | painful | occipital | right | 20 | -3.35 | 2.53 |
| TS | enhanced | painful | occipital | left | 20 | -3.52 | 2.58 |
| TS | enhanced | painful | occipital | middle | 20 | -2.48 | 2.45 |
| TS | enhanced | painful | occipital | right | 20 | -3.40 | 2.65 |

**Table S10**: Anova with factors Group, Sensitivity, Action, Region and Laterality:

| Effect | DFn | DFd | F | p | p<.05 | ges |
|---|---|---|---|---|---|---|
| Group | 1 | 41 | 1.06 | 0.310 | | 0.02 |
| Sensitivity | 1 | 41 | 0.04 | 0.848 | | 0.00 |
| Action | 1 | 41 | 9.47 | 0.004 | * | 0.01 |
| Region | 1 | 41 | 13.84 | 0.001 | * | 0.04 |
| Laterality | 2 | 82 | 8.51 | 0.000 | * | 0.01 |
| Group:Sensitivity | 1 | 41 | 0.84 | 0.364 | | 0.00 |
| Group:Action | 1 | 41 | 0.09 | 0.764 | | 0.00 |
| Group:Region | 1 | 41 | 2.31 | 0.136 | | 0.01 |
| Group:Laterality | 2 | 82 | 0.15 | 0.865 | | 0.00 |
| Sensitivity:Action | 1 | 41 | 0.27 | 0.608 | | 0.00 |
| Sensitivity:Region | 1 | 41 | 0.36 | 0.552 | | 0.00 |
| Action:Region | 1 | 41 | 4.56 | 0.039 | * | 0.00 |
| Sensitivity:Laterality | 2 | 82 | 1.59 | 0.211 | | 0.00 |
| Action:Laterality | 2 | 82 | 0.75 | 0.476 | | 0.00 |
| Region:Laterality | 2 | 82 | 8.68 | 0.000 | * | 0.01 |
| Group:Sensitivity:Action | 1 | 41 | 5.31 | 0.026 | * | 0.00 |
| Group:Sensitivity:Region | 1 | 41 | 0.95 | 0.336 | | 0.00 |
| Group:Action:Region | 1 | 41 | 0.27 | 0.605 | | 0.00 |
| Group:Sensitivity:Laterality | 2 | 82 | 0.81 | 0.446 | | 0.00 |
| Group:Action:Laterality | 2 | 82 | 2.18 | 0.120 | | 0.00 |
| Group:Region:Laterality | 2 | 82 | 0.68 | 0.507 | | 0.00 |
| Sensitivity:Action:Region | 1 | 41 | 3.84 | 0.057 | | 0.00 |
| Sensitivity:Action:Laterality | 2 | 82 | 1.00 | 0.372 | | 0.00 |
| Sensitivity:Region:Laterality | 2 | 82 | 0.60 | 0.549 | | 0.00 |
| Action:Region:Laterality | 2 | 82 | 0.92 | 0.403 | | 0.00 |
| Group:Sensitivity:Action:Region | 1 | 41 | 0.30 | 0.587 | | 0.00 |
| Group:Sensitivity:Action:Laterality | 2 | 82 | 2.49 | 0.089 | | 0.00 |
| Group:Sensitivity:Region:Laterality | 2 | 82 | 1.58 | 0.213 | | 0.00 |
| Group:Action:Region:Laterality | 2 | 82 | 0.28 | 0.756 | | 0.00 |
| Sensitivity:Action:Region:Laterality | 2 | 82 | 1.14 | 0.325 | | 0.00 |
| Group:Sensitivity:Action:Region:Laterality | 2 | 82 | 3.37 | 0.039 | * | 0.00 |

**Table S11**: Anova results grouped by Region:

| Region | Effect | DFn | DFd | F | p | p<.05 | ges |
|---|---|---|---|---|---|---|---|
| central | Group | 1 | 41 | 3.36 | 0.074 | | 0.06 |
| central | Action | 1 | 41 | 2.87 | 0.098 | | 0.00 |
| central | Sensitivity | 1 | 41 | 0.22 | 0.641 | | 0.00 |

| | | | | | | | |
|---|---|---|---|---|---|---|---|
| central | Laterality | 2 | 82 | 0.95 | 0.392 | | 0.00 |
| central | Group:Action | 1 | 41 | 0.00 | 0.981 | | 0.00 |
| central | Group:Sensitivity | 1 | 41 | 0.26 | 0.616 | | 0.00 |
| central | Group:Laterality | 2 | 82 | 1.03 | 0.361 | | 0.00 |
| central | Action:Sensitivity | 1 | 41 | 0.52 | 0.475 | | 0.00 |
| central | Action:Laterality | 2 | 82 | 0.00 | 0.997 | | 0.00 |
| central | Sensitivity:Laterality | 2 | 82 | 0.09 | 0.913 | | 0.00 |
| central | Group:Action:Sensitivity | 1 | 41 | 6.52 | 0.014 | * | 0.00 |
| central | Group:Action:Laterality | 2 | 82 | 2.10 | 0.129 | | 0.00 |
| central | Group:Sensitivity:Laterality | 2 | 82 | 0.13 | 0.876 | | 0.00 |
| central | Action:Sensitivity:Laterality | 2 | 82 | 2.02 | 0.140 | | 0.00 |
| central | Group:Action:Sensitivity:Laterality | 2 | 82 | 1.82 | 0.169 | | 0.00 |
| occipital | Group | 1 | 43 | 0.00 | 0.948 | | 0.00 |
| occipital | Action | 1 | 43 | 13.80 | 0.001 | * | 0.01 |
| occipital | Sensitivity | 1 | 43 | 0.12 | 0.735 | | 0.00 |
| occipital | Laterality | 2 | 86 | 13.73 | 0.000 | * | 0.02 |
| occipital | Group:Action | 1 | 43 | 0.61 | 0.440 | | 0.00 |
| occipital | Group:Sensitivity | 1 | 43 | 2.09 | 0.156 | | 0.00 |
| occipital | Group:Laterality | 2 | 86 | 0.52 | 0.598 | | 0.00 |
| occipital | Action:Sensitivity | 1 | 43 | 2.73 | 0.106 | | 0.00 |
| occipital | Action:Laterality | 2 | 86 | 1.23 | 0.299 | | 0.00 |
| occipital | Sensitivity:Laterality | 2 | 86 | 1.36 | 0.262 | | 0.00 |
| occipital | Group:Action:Sensitivity | 1 | 43 | 3.14 | 0.083 | | 0.00 |
| occipital | Group:Action:Laterality | 2 | 86 | 1.88 | 0.160 | | 0.00 |
| occipital | Group:Sensitivity:Laterality | 2 | 86 | 1.57 | 0.215 | | 0.00 |
| occipital | Action:Sensitivity:Laterality | 2 | 86 | 0.05 | 0.947 | | 0.00 |
| occipital | Group:Action:Sensitivity:Laterality | 2 | 86 | 2.81 | 0.066 | | 0.00 |

**Table S12:** Anova results for central regions, grouped by Group:

| Group | Effect | DFn | DFd | F | p | p<.05 | ges |
|---|---|---|---|---|---|---|---|
| HC | Action | 1 | 24 | 1.56 | 0.224 | | 0.00 |
| HC | Sensitivity | 1 | 24 | 0.67 | 0.421 | | 0.00 |
| HC | Laterality | 2 | 48 | 0.23 | 0.798 | | 0.00 |
| HC | Action:Sensitivity | 1 | 24 | 5.11 | 0.033 | * | 0.00 |
| HC | Action:Laterality | 2 | 48 | 1.19 | 0.314 | | 0.00 |
| HC | Sensitivity:Laterality | 2 | 48 | 0.24 | 0.785 | | 0.00 |
| HC | Action:Sensitivity:Laterality | 2 | 48 | 2.59 | 0.085 | | 0.00 |
| TS | Action | 1 | 17 | 1.48 | 0.241 | | 0.00 |
| TS | Sensitivity | 1 | 17 | 0.00 | 0.983 | | 0.00 |
| TS | Laterality | 2 | 34 | 2.33 | 0.113 | | 0.01 |
| TS | Action:Sensitivity | 1 | 17 | 2.24 | 0.153 | | 0.00 |
| TS | Action:Laterality | 1.5 | 25.53 | 0.95 | 0.377 | | 0.00 |
| TS | Sensitivity:Laterality | 2 | 34 | 0.00 | 0.998 | | 0.00 |
| TS | Action:Sensitivity:Laterality | 2 | 34 | 1.44 | 0.252 | | 0.00 |

**Table S13**: Pairwise comparisons of action per sensitivity and group:

| Group | Sensitivity | variable | group1 | group2 | n1 | n2 | p |
|---|---|---|---|---|---|---|---|
| HC | normal | value | neutral | painful | 75 | 75 | 0.165 |
| HC | enhanced | value | neutral | painful | 75 | 75 | 0.917 |
| TS | normal | value | neutral | painful | 58 | 59 | 0.695 |
| TS | enhanced | value | neutral | painful | 60 | 60 | 0.399 |

### Skin Conductance Analyses

#### SCR recording

Skin conductance response (SCR) was recorded using two electrodes attached to the medial phalanges of the index and middle fingers of the left hand. The online reference was placed on the left forearm and recorded with the same amplifier (BrainAmp ExG. BrainProdcts GmbH). As SCR data were measured simultaneously with EEG data, the sampling rate was 500 Hz, and an online high-pass filter of 0.016 Hz as well as a notch filter of 50 Hz were applied.

#### SCR data analysis

Analysis of skin conductance was performed using the MATLAB® toolbox *Ledalab* (Benedek & Kaernbach, 2010). The signal was downsampled to 25 Hz and then smoothed with a Gaussian low-pass filter. Obvious artifacts were removed manually. Then, the signal was deconvolved using a continuous decomposition analysis (CDA) with four optimizations. This separates the signal into continuous traces of tonic and phasic activity. We proceeded to analyze the phasic driver component, by extracting averaged event-related responses via area under the curve, between 1 to 4 seconds after stimulus onset. Event-related phasic SCR were range normalized and log transformed across all trials within each subject. This ensured that individual variability in the signal magnitude would not bias further analyses. SCR per trial were then averaged within conditions. One participant from each group was excluded, as the data quality did not allow measurements of any SCR. Thus, analyses of SCR data were conducted on n = 20 TS patients and n = 24 healthy controls. SCR were analyzed similarly to the EEG data with a mixed ANOVA with the factors group, action and sensitivity. For correlation analyses, difference scores were calculated analogous to the behavioral and EEG data.

#### SCR results

Analysis of the skin conductance response revealed a significant interaction of group, action and sensitivity ($F(1.42) = 5.40$. $p = .025$. $\eta^2 = .01$; Fig. S1). ANOVAs per group revealed a main effect of action in the TS group ($F(1.19) = 4.39$. $p = .050$. $\eta^2 = .01$), with participants showing an enhanced skin conductance response after watching painful actions. For the healthy controls we found an interaction of action and sensitivity ($F(1.23) = 6.24$. $p = .020$. $\eta^2 = .01$), with nominally stronger responses after painful pictures in the enhanced sensitivity condition. Pairwise comparisons were not significant.

#### Discussion of SCR results

Physiologically, increased distress due to increased sharing should have been visible in a stronger skin conductance response in the TS group. However, we observed no such group differences. In fact, the results suggested a slightly reduced modulation of skin conductance response by the sensitivity manipulation in the TS group. Previous research has linked the magnitude of SCR in response to witnessing others in pain with prosocial behavior (Hein et al., 2011) and implicit race bias (Forgiarini et al., 2011), where higher SCRs corresponded to increased empathy and prosocial behavior. In conclusion, our findings may suggest a slight reduction in empathic responses among individuals with

TS compared to the control group. However, it is important to acknowledge that the stimuli in our paradigm were presented quite fast, which may have caused overlapping of skin conductance responses. In contrast, previous studies examining SCR in empathy for pain paradigms (Forgiarini et al., 2011; Hein et al., 2011) used longer inter-trial intervals, and the painful stimuli used were more salient, such as observing hands being pricked by needles or individuals receiving electrical shocks. Overall, it is possible that our experimental design might have caused us to miss subtle group differences. Nonetheless, there was no evidence indicating an increased physiological response to painful situations in the TS group.

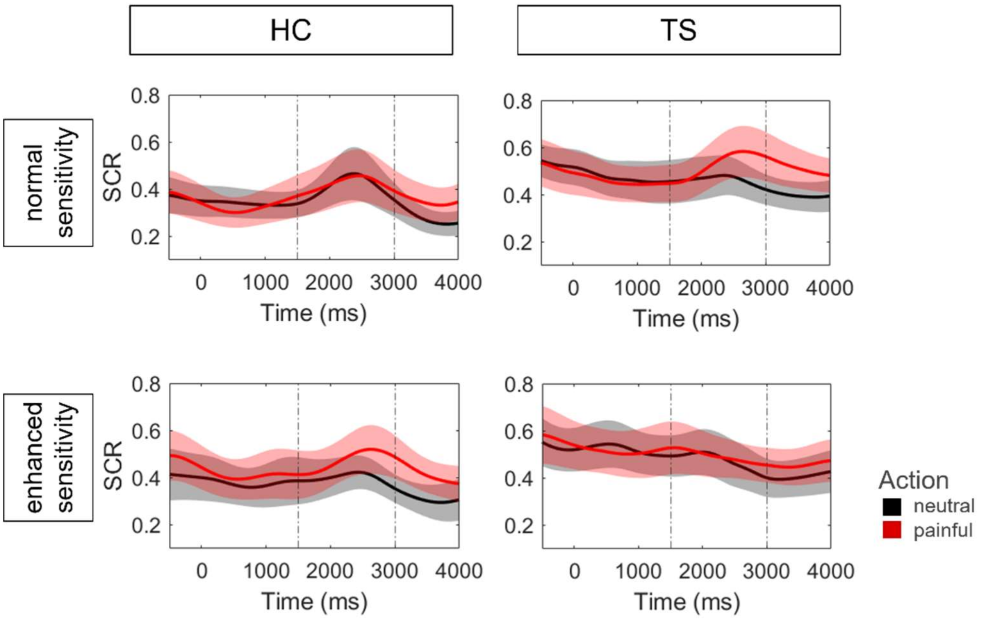


**Figure S1.** Skin conductance response in empathy for pain paradigm. Plotted are the time courses of the adjusted phasic driver for the painful (red) and neutral (black) pictures, separately for the controls (HC, left columns) and Tourette patients (TS, right columns) and for normal and enhanced sensitivity. Time-point 0 refers to the picture onset. Statistical analysis is based on our window of interest (1500 - 3000 ms).
